# The Minimal Dataset for Cancer of the 1+Million Genomes Initiative

**DOI:** 10.1101/2023.10.07.561259

**Authors:** Michela Riba, Cinzia Sala, Aedin Culhane, Åsmund Flobak, Attila Patocs, Kjetil Boye, Karla Plevova, Šárka Pospíšilová, Giorgia Gandolfi, Marco J Morelli, Gabriele Bucci, Anders Edsjö, Ulrik Lassen, Fátima Al-Shahrour, Nuria Lopez-Bigas, Randi Hovland, Edwin Cuppen, Alfonso Valencia, Helene Antoine-Poirel, Richard Rosenquist Brandell, Serena Scollen, Juan Arenas Marquez, Jeroen Belien, Arcangela De Nicolo, Ruggero De Maria, David Torrents, Giovanni Tonon

## Abstract

For a real impact on healthcare, precision cancer medicine requires accessibility and interoperability of clinical and genomic data across centres and countries. Due to the heterogeneous digitization in Europe and worldwide, the definition of models for standardised data collection and usability becomes mandatory if countries want to work together on this mission. The European Union 1+Million Genomes (1+MG) initiative, supported by the Horizon 2020 Beyond 1 Million Genomes project, aims at outlining data models, guidance, best practices, and technical infrastructures for transnational access to sequenced genomes, including cancer genomes. Within the framework of the cancer-focused Working Group 9, we developed the 1+MG-Minimal Dataset for Cancer (1+MG-MDC)–a data model encompassing 140 items and organized in eight conceptual domains for the collection of cancer-related clinical information and genomics metadata. The 1+MG-MDC, which results from a multidisciplinary effort, leverages pre-existing models and emphasizes the annotation and traceability of multiple aspects relevant to the complex longitudinal path of the cancer disease and its treatment. We strived to make the 1+MG-MDC easy to adopt, yet comprehensive, addressing the needs of both clinicians and researchers. We will periodically revise and update it to ensure it remains fit for purpose. We propose the 1+MG-MDC as a model to create homogeneous databases, which would, in turn, guide discussions on clinical and genomic features with prognostic or therapeutic value and foster real-world data research.

## Introduction

Genome sequencing plays a central role in precision medicine. Detailed knowledge of genomic variation is essential to elucidate the molecular bases of diseases, to identify genetic predisposition, to refine diagnosis, to guide therapeutic choices and follow-up, and to identify prognostic and predictive signatures. This is especially true for cancer – a disease for which diagnosis relies more and more on genetic biomarkers and treatment is increasingly tailored to the genetic alterations present in malignant cells.

It is becoming increasingly evident that for a tangible impact on healthcare, data need to be accessible and interoperable at a scale that has not been achieved yet in Europe. Nor, for that matter, in most countries worldwide. The sharing of clinical and genomic data is a matter of lively debates within the European Union (EU) scientific community ^1,2^. Several initiatives have been launched to tackle the ethical, legal, political, and technical challenges that need to be addressed both at the country and the EU level ^3^. Clinical and genomic data sharing will require multi-institutional and multi-national collaboration ^4^ and will provide a data platform of unprecedented size that will empower next generation precision medicine research. Federated learning, relying on common interoperable data standards, infrastructures, and platforms, will be especially valuable.

The 1+ Million Genomes (1+MG) initiative was launched in 2018 within the framework of the EU agenda for the Digital Transformation of Health and Care (https://digital-strategy.ec.europa.eu/en/policies/ehealth). Its mission is to enable clinical and genomic data federation across Europe, by establishing infrastructure, legal guidance, and best practices, to allow secure cross-border access to more than one million sequenced genomes, including cancer genomes (https://digital-strategy.ec.europa.eu/en/policies/1-million-genomes).

The 1+MG activities have been organised through 12 Working Groups (WGs), including WG3 on *Common standards and minimal dataset for clinical and phenotypic data*, and five WGs focusing on use cases, specifically, *Rare diseases* (WG8), *Cancer* (WG9), *Common and complex diseases* (WG10), *Infectious diseases* (WG11), *and Population genomics* (WG12) (https://b1mg-project.eu/about/how-organised/operational-group). The 1+MG initiative was supported by the Horizon 2020 Beyond 1 Million Genomes (B1MG) project, structured in seven work packages, covering a variety of tasks, from stakeholder engagement to technical infrastructure to precision medicine delivery and impact (https://b1mg-project.eu/). To date, 25 EU countries, along with Norway and the United Kingdom, have joined the initiative. In November 2022, the Genomic Data Infrastructure (GDI) project (https://gdi.onemilliongenomes.eu), funded by the EU Digital Europe programme, was launched. GDI builds upon B1MG and aims at realising the promise of 1+MG by establishing a European data infrastructure for secure cross-border access to genomic data.

The 1+MG WG9 is working towards actualising federated data query and access, whereby first a search for genomic and matched clinical data meeting specific criteria is run to identify which European centres have the information available and then access to these data is requested. In the context of the WG9 activities and discussions, the need for a Common Data Model (CDM), which would capture essential genomic and clinical information for each cancer patient, emerged as a priority. Indeed, cancer presents peculiarities that set it apart from other conditions. For instance, rare genetic diseases are typically driven by germline DNA changes and sequencing for diagnostic purposes is performed only once. Instead, cancer may be characterized by various types of (both germline and somatic) genetic events, often endowed with very intricate genomic changes. Coexistence of different types of alterations (*e*.*g*., single nucleotide variants, small and large deletions and insertions, complex structural variants), which can differently impact on genome function, complicates the interpretation of the genomic landscape. Moreover, oftentimes a cancer patient receives multiple lines of treatment due to resistance, disease recurrence, or metastases–events that may be associated with novel somatic genetic changes. Furthermore, because multiple targeted treatments are nowadays available, to guide cancer precision treatment, patients may have multiple genetic tests, on different sequencing platforms at different times, and these multi-temporal data generate complex follow-up information. All these features require *ad hoc* data models, which cannot be easily adopted from other settings. A CDM would therefore serve as a reference scheme that provides guidance, based on specific standards, for observational data collection, thus ensuring uniform and harmonised structure and content of the modelled databases and facilitating subsequent (single centre, centralised, or federated) data analyses. Agreement on a “core” set of data to be collected from cancer patients is prerequisite to achieving comparability and, ultimately, interoperability, whereas excessive discretionary power could hinder wider searches and analyses, hampering potential improvements in patient care. A CDM will also be helpful for legislative purposes, clarifying what information needs to be shared for research and development and health care programs.

For the definition of a CDM, in addition to establishing a common set of data items relevant to specific semantic domains (*e*.*g*., demographics, disease-related features, therapy), it is paramount to specify how those data should be reported, to maximize the use of standards and ensure interoperability, and to limit the presence of unstructured free text. Standard value sets (*i*.*e*., specific values, terms, or codes for data element description) allow population of a database in a structured manner. Data dictionaries assist with standardization, providing coded medical concepts. They can be either comprehensive (*i*.*e*., encompassing various domains) or domain-specific (*i*.*e*., referring to specific data such as disease classification and drugs) (**Supplementary Table 1**). Some dictionaries are freely accessible; others require a license and/or access fee.

Currently, the largest available dictionaries are the so-called “metathesauri”– vocabulary databases, often multilingual, built from several thesauri or domain-specific dictionaries. The Unified Medical Language System (UMLS) metathesaurus by the National Institutes of Health is a multilingual dictionary encompassing biomedical and health-related definitions. Assembled from several different electronic standard vocabularies, it contains more than five million terms, each with a unique identifier ^6^. Other available dictionaries cover concepts relating to specific domains. For instance, the HemOnc database (https://hemonc.org/) ^7,8^, a structured wiki of hematology and oncology drugs and treatments, provides coded cancer treatments including drug combinations and defined regimens of administration. The Observational Health Data Sciences and Informatics (OHDSI) Observational Medical Outcomes Partnership (OMOP) CDM is an open community data standard providing both a common data model (OMOP CDM) and a vocabulary repository (ATHENA, https://athena.ohdsi.org/) that maps over 60 medical vocabularies, including Systematic Nomenclature of Medicine -Clinical Terms (SNOMED-CT), Logical Observation Identifiers Names and Codes (LOINC), and International Classification of Diseases (ICD10), to be used with the OMOP CDM. Other international standards referring to more general categories may serve as a source of value sets not specific to medicine-, science-, and biology-related semantics (*e*.*g*., date standards refer to ISO specifications).

A faithful and comprehensive description of a cancer patient encompasses several layers of clinical and genomic information. Cancer clinical history unfolds longitudinally, and may include different disease stages (primary cancer, relapse, additional recurrences/transformation) and treatments (*e*.*g*., cycles of therapy, surgical procedures, radiotherapy, various time points and methods of response assessment), thus resulting in a complex and nested picture. From a genetic standpoint, cancer genome assessment should include both germline sequence variants and somatic changes, which can predict the clinical behaviour and prognosis and determine response or resistance to certain treatments, in addition to discriminating diagnosis and predicting potential cancer risk for blood relatives. Because somatic changes may evolve over time or following treatment and may vary by cellular sources (*e*.*g*., local or regional metastatic site, liquid biopsy), genomic data can have multiple layers. Current databases collecting cancer genomic data generally lack some dimensions since they were designed each to capture a specific aspect of the cancer disease. Increasingly an individual may have multiple genomics assays, targeted sequencing on comprehensive gene panel, exome, whole genome, single cell sequencing in addition to immuno-profiling, proteomics or other omics approaches.

The general aim of the 1+MG initiative and the specific focus of the WG9 on cancer prompted us to define a Minimal Dataset for Cancer (1+MG-MDC), which we envisioned as a model that would be both minimal (*i*.*e*., include a reasonably limited number of items that are easy to collect for health professionals, who usually fill in this information within an already severely constrained schedule) and yet comprehensive, encompassing structured data that are useful for both clinicians and researchers.

## Methods

To define the key data domains and the corresponding items to be included in our 1+MG-MDC, we employed a strategy encompassing two surveys and several rounds of discussions and involving all WG9 members or *ad hoc* focus expert groups (**Figure 1**).

**Figure 1.**
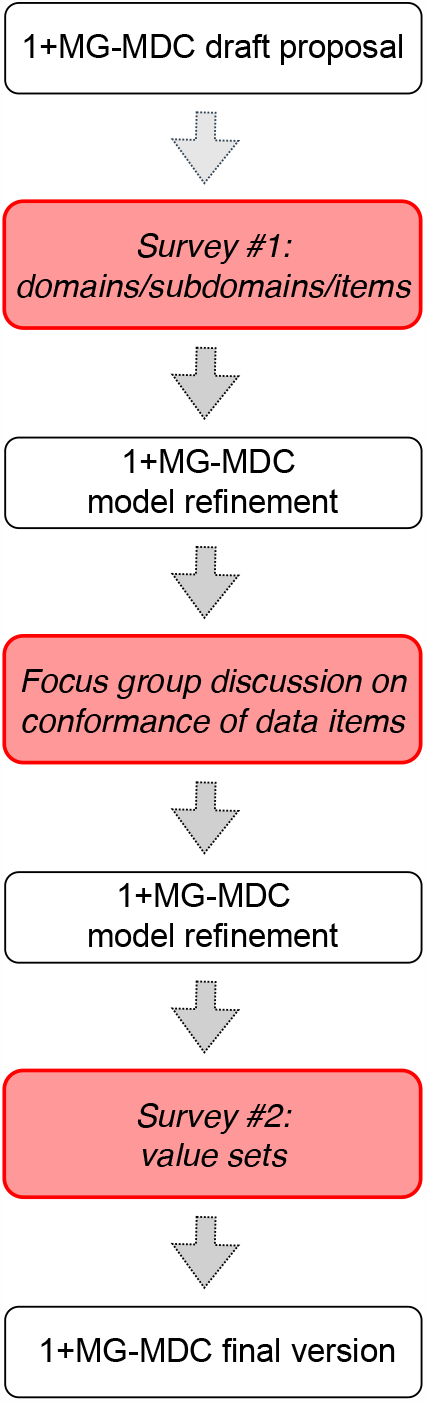
Strategy to develop the 1+MG-MDC. Flowchart summarizing the strategy employed to develop the 1+MG-MDC.

After systematic revision of the data models currently used for clinical data collections and within bioinformatics initiatives (see **Table 1**), we compiled 13 tables, each corresponding to an operational domain (namely, *Submitter, Patient, Sequence, Sample, Disease Cancer, Disease Comorbidity, Treatment, Treatment Cancer Surgery, Treatment Cancer Chemotherapy, Treatment Cancer Radiation Therapy, Treatment Cancer Hormonal Therapy and Biological Therapy, Environmental Exposure, Biomarker*) and including related items (**Table 2**). We then invited the 1+MG WG9 members to participate in a general survey (Survey #1 -defining the data domains and associated items) via a web platform. For each of the proposed items (n=145), all participants were asked to specify whether they concurred with inclusion or not, providing comments and suggestions. This step was followed by a collegial review and discussion of the results. An *ad hoc* team of experts was established to tackle the most challenging items, leading to the removal of 48 (18 from *Patient*, 14 from *Environmental Exposure*, one from *Treatment Cancer Chemotherapy*, 11 from *Treatment Cancer Hormonal Therapy and Biological Therapy*, two from *Biomarker*, one from *Sequence*, and one from *Treatment*) and addition of 43 (38 relating to *Sample*, two to *Patient*, one to *Environmental Exposure*, one to *Treatment Cancer Hormonal Therapy and Biological Therapy*, one to *Treatment Cancer Surgery*) items.

**Table 1.**
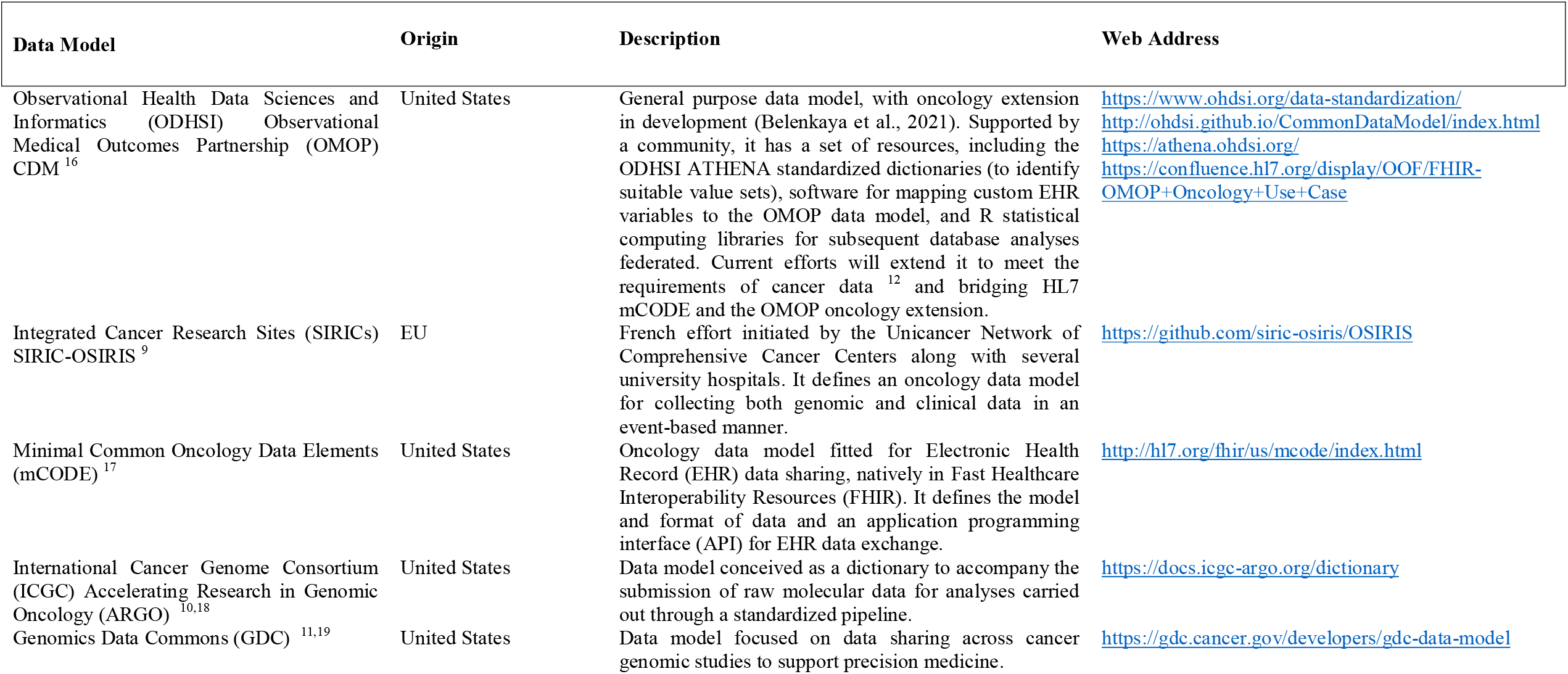
Data models used as a reference to build the Minimal Dataset for Cancer.

**Table 2.**
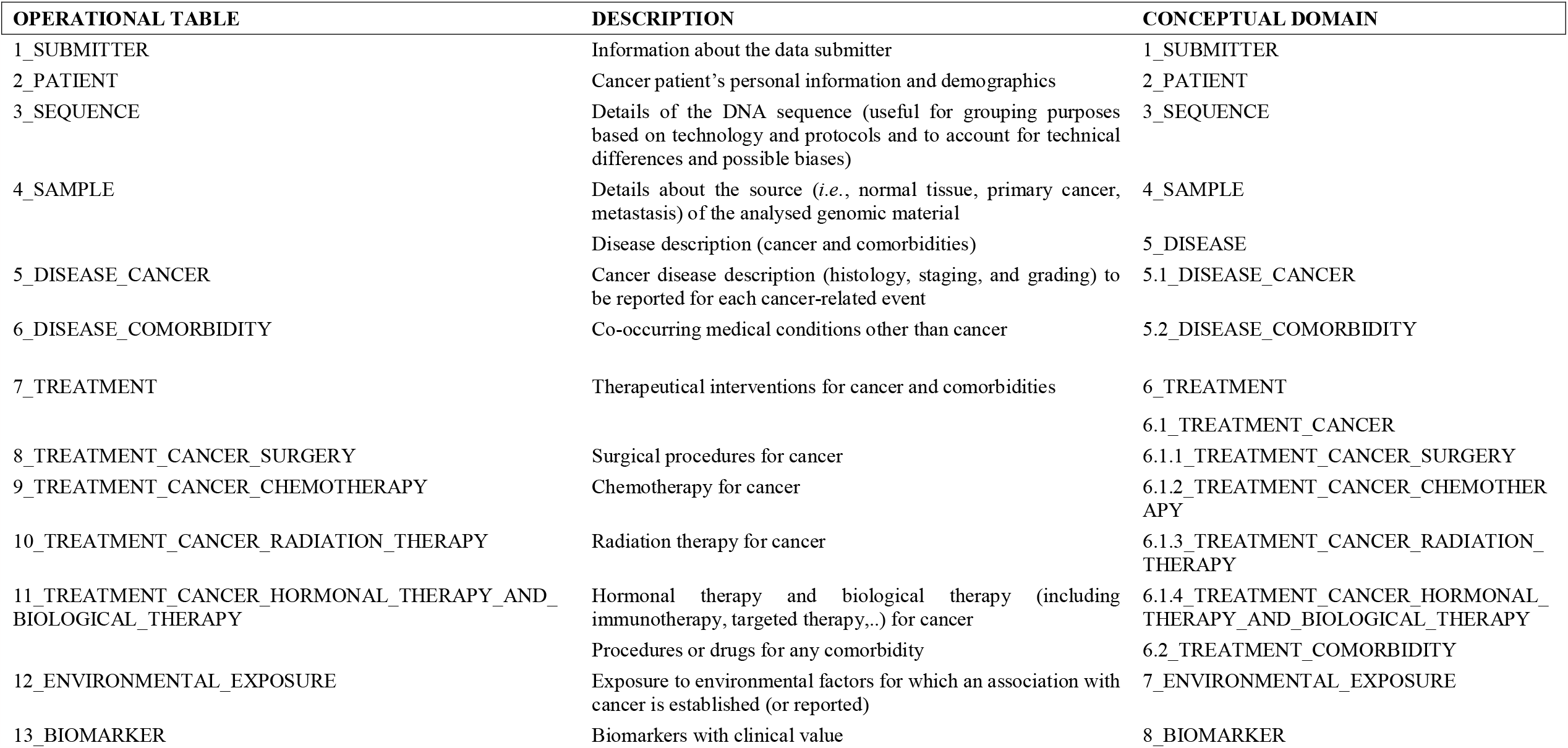
List of the eight conceptual domains (and corresponding 13 operational tables) covered by the Minimal Dataset for Cancer.

Next, a focus group convened to discuss item conformance. Based on a balanced agreement between various stakeholders, including clinicians and researchers, the items were assigned to three different bins: *mandatory (i*.*e*., core items such as demographics, age, sex at birth, sequencing protocols, cancer pathology, etc.), *recommended* (*i*.*e*., sample processing or cancer treatment details, which are highly valuable for research, yet often cumbersome for clinicians to fill and/or demanding for researchers to collect systematically and usually not required in databases of primary cancer genomics collections) or *optional* (*i*.*e*., environmental exposures or treatment information for comorbidities, which are not strictly connected with the goal of data collection, yet valuable for collateral analyses).

A second survey (Survey #2 -defining the value sets), included the updated and refined list of 140 items relating to the 13 operational tables, was administered aiming at a majority consensus by WG9 members with expertise in medical oncology, cancer biology, genetics, and bioinformatics on the value sets, approved by different professionals, to be used for the compilation of the forms collecting data, which could be potentially built from the data model. For this purpose, the use of free-of-charge data dictionaries was prioritized to ensure inclusiveness and equality. The feedback received was further discussed and persistent disagreement resolved based on a majority consensus The final 1+MG-MDC, with the proposed values, is shown as **Supplementary File 1**, which also includes references to the published data models used as a source or reference, depending on complete or partial (if the definition and/or proposed value set slightly differed) correspondence with their items.

## Results

Using a stepwise approach and building upon existing data models, we defined and herein propose the 1+MG-MDC, which aims to capture the complex evolution of cancer, by enabling the collection of data obtained at different times of its longitudinal path (*e*.*g*., clinical information associated with genomic data, from more than one cancer sample from the same patient). Our strategy offers the advantage of diminishing the load of field-specific items while highlighting those collegially supported by the various professionals involved in the modelling process, including clinicians upon whom most of the burden for data collection lies. By cross-referencing the 1+MG-MDC with existing data models (and related items), we leveraged current knowledge and experience while moving towards data collections encompassing comprehensive longitudinal sequencing metadata and clinical information.

The 13 tables, collated in eight conceptual domains (**Table 2**), reflect the needs of clinical and genomics cancer data collection and aim to balance data completeness and the limited time that could be realistically allocated to data compilation. The identification of a set of core items, without which a new submission of clinical and genomic data would not be possible, guarantees that fundamental information is included in all collections.

Starting from the initially proposed 145 items, through a process of multidisciplinary plenary as well as small focus group discussions, we obtained a final list of 140 (37 *mandatory*, 40 *recommended* and 63 *optional*) items relevant to the eight conceptual domains. Specifically, the 1+MG-MDC includes 13 items associated with the *Submitter* domain, five with the *Patient*, nine with the *Sequence*, 45 with the *Sample*, 16 with the *Disease* (13 with the *Disease Cancer* and 3 with *Disease Comorbidity*, respectively), 44 with the *Treatment* (specifically, two with *Treatment Cancer*; two with *Treatment Cancer Surgery*, 13 with *Treatment Cancer Chemotherapy*, 14 with the *Treatment Cancer Radiation Therapy*, ten with *Treatment Cancer Hormonal Therapy and Biological Therapy*, three with *Treatment Comorbidity*), five with the *Environmental Exposure*, and three with the *Biomarker* domains (**Figure 2**).

**Figure 2.**
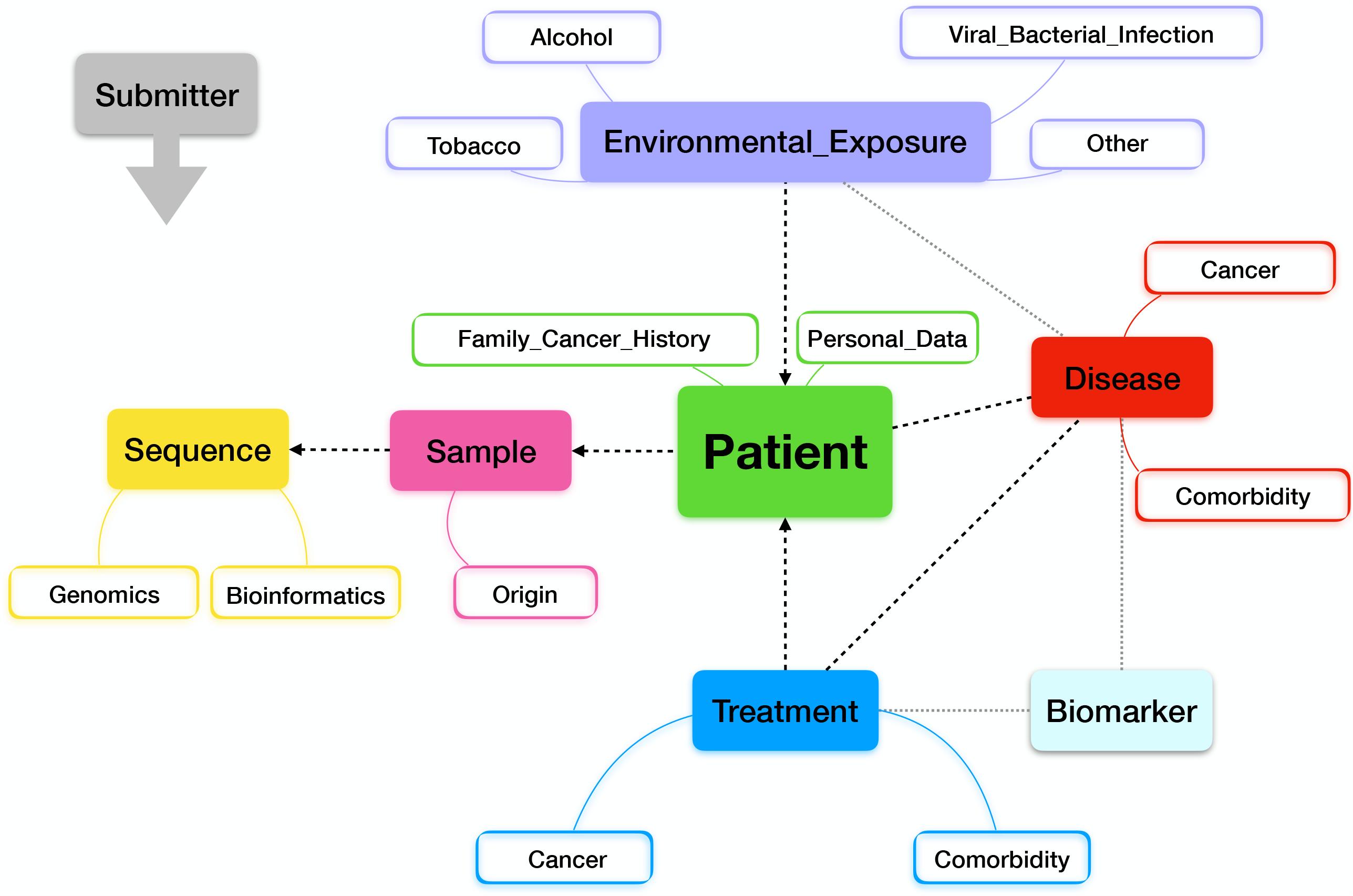
The 1+MG-MDC structure. Graph illustrating the 1+MG-MDC structure, comprising eight conceptual domains and related subdomains.

Information regarding the *Submitter* allows data stratification based on participating centre and/or country. The information collected in the *Patient* domain (demographics) encompasses a list of items suitable for all countries. We did not include items that cannot be collected in some EU member states (*e*.*g*., ethnicity, for legal reasons).

One of the key features of the 1+MG-MDC is the information about the *Sequence* domain. It was agreed that it should contain metadata useful to classify the sequence, including file names with either raw or pre-processed extension (*e*.*g*., “.fastq.gz” and “.vcf”, respectively), genomic preparation protocols, sequencing platforms (*i*.*e*., single gene, gene panels, exome or whole-genome sequences), and bioinformatics pipelines used for data processing, generation (made available in version-tracking repositories, for example the ELIXIR Workflow Hub, https://workflowhub.eu/ Goble et al., 2021), and interpretation. The decision to acquire many sequence types was based on the anticipation that a future database, built based on the 1+MG-MDC, should be accessible and easy to use by multiple centres regardless of their technological development and the local availability of dedicated personnel for data management and database filling.

As per discussions and agreement within the WG9, we chose to be accurate with regards to the *Sample* domain, which comprises a total of 45 items, to account for the various sources the sequences could be derived from, ranging from primary solid tumours or haematological malignancies to metastatic specimens and liquid biopsy. *Disease Cancer* includes information about staging and grading (eventually with different classification protocols applying to specific malignancies). Data regarding co-occurring pathologies (*Disease Comorbidity*) are also collected. The *Treatment* domain details surgical procedures or pharmacological treatments for comorbidities and includes several cancer treatment-specific subdomain categories. Indeed, owing to the inherent complexity of the cancer therapy path, adequate space has been given to the specification of the different treatment types, ranging from surgery to various therapy regimens, specifying their different purposes (*e*.*g*., neo-adjuvant versus adjuvant chemotherapy) and possible multiple lines, reporting dates of treatment initiation/end (*Treatment Cancer Surgery, Treatment Cancer Chemotherapy, Treatment Cancer Radiation Therapy, Treatment Cancer Hormonal Therapy and Biological Therapy* subdomain categories). These parameters could serve to extract outcomes, such as time to next treatment, which is useful in real-world evidence research. Adverse events and outcome are also included as optional and recommended items, respectively. The simple identification of the code of an active substance does not allow proper description of a complex set of drugs administered through numerous cycles; we addressed the intricacy associated with a standardized classification of cancer therapies by identifying a suitable vocabulary of value-sets and avoiding the use of unstructured free text.

The *Environmental Exposure* domain, structured for tobacco, alcohol and viral/bacterial infection, also allows the inclusion of other factors (*e*.*g*., diet, professional exposures, etc.). Known *Biomarkers* associated to diagnosis, prognosis, or treatment (*e*.*g*., an actionable sequence variant in a single gene) or measurable residual disease can be included in the corresponding domain.

## Discussion

Standardised data collections are a prerequisite for making data querying, and eventually data sharing, useful and valuable for clinical decision making. Differences exist across European countries and even among centres within the same country, which reflects heterogeneity in the level of digitalisation of medical information. These range from structured and centralized solutions to centre- or project-specific databases with a high degree of unstructured data often collected as free text clinical notes. Defining minimal standards for data collection is key to obtaining interoperable high-quality databases that could be used in real-world evidence cancer research and contribute to advancing precision treatment options. Without this standardisation, it will be more difficult to perform translational research and clinical studies, slowing down the advances in precision cancer medicine.

Under the 1+MG-WG9 umbrella, we conceived and developed the 1+MG-MDC to enable the collection of cancer genomes in Europe as well as associated clinical data. Capitalising upon the most authoritative and available data models (**Supplementary Table 1**) and leveraging on their strengths, we constructed a comprehensive yet limited data standard. As an example, we took into account the event-based structure for an accurate definition of the cancer longitudinal path from OSIRIS ^9^; the cancer genome focus from ICGC-ARGO ^10^; the detailed description of the bioinformatics procedures from GDC ^11^; and the comprehensive medical vocabulary curation from ODHSI-OMOP ^12^.

Today, cancer studies collect different, only partially overlapping, sets of variables, depending on their specific aims. The 1+MG-MDC encompasses items pertaining to eight semantic domains and aims at fostering the collection of clinical information and genomic data from cancer patients across Europe for both research and clinical use. Notably, the 1+MG-MDC accounts for the possibility that genomics and clinical data are used beyond clinical trial purposes in a real-world evidence perspective ^13^. We ensured that the 1+MG-MDC could be easy to use for both clinicians and researchers, as well as by new emerging professional categories dealing with real-world data collection, such as physician scientists, clinical information engineers, research nurses, data managers, epidemiologists, and health care providers.

The 1+MG-MDC addresses European-specific needs to allow the collection of a large amount of cancer genome and associated clinical data contributed by many centres across Europe regardless of their medical digitization and technology development. The introduction of mandatory and recommended items should theoretically ensure the collection of “difficult to have” data–a major strength of the 1+MG-MDC, which can include details relevant to sequence, samples, and therapies at different points of the cancer disease path, while still maintaining the number of the mandatory items within a workable limit of less than 40.

Notably, for the 1+MG-MDC, we used free-of-charge dictionaries to ensure wide utilization since the use of a comprehensive, freely accessible dictionary ensures a unified source of value sets. A possible exception is represented by the largest available multilingual health dictionary, Systemized Nomenclature of Medicine Clinical Terms (SNOMED CT), which can currently be accessed in most EU countries free of charge. Indeed, the EU Commission is supporting the use of SNOMED CT through direct grants to Member States to cover joining and membership fees. In case, since back-compatibility for most dictionaries used to create the 1+MG-MDC is expected, the future alignment with SNOMED would be pursuable.

We propose the 1+MG-MDC as a blueprint to construct databases for standardized data collection, overcoming currently existing hurdles to collaborative trans-national genomic studies and maximizing the interoperability among different cancer centres throughout Europe and beyond. We hope that the 1+MG-MDC may support molecular tumour boards ^14,15 1415^and contribute to advancement in precision cancer care. The real test bench will be the use by clinicians, researchers, and other healthcare providers, who could corroborate its value, suggest further refinements, and advise on updates for alignment with evolving classifications.

## Conclusions

We propose the 1+MG-MDC, a data model for the collection of genomic and associated clinical data from cancer patients. The 1+MG-MDC was built leveraging upon the existing data models for cancer genomics in Europe and beyond and it is the result of a multidisciplinary effort that involved oncologists, haematologists, pathologists, geneticists, surgeons, molecular biologists, bioinformaticians, and medical informaticians. It includes 140 data items organized into semantically coherent domains, and it will serve as a guide to construct databases for the collection of clinical and genomic information. Conceived and developed within the framework of the EU 1+MG initiative and EU-funded Horizon 2020 B1MG project, the 1+MG-MDC could potentially assist molecular tumour boards towards a shared common language for collegial discussions of clinical and genomics features important to determine the best pathway of care for a cancer patient. Promoting the use of standards, the 1+MG-MDC will foster trans-national collaborations. It will also serve as an exemplar for future integration of other omics layers.

## Figure Legends

**Supplementary File 1. The 1+MG-MDC** Columns from left to right: *Item number*, integer indicating the data item; *Domain*, indicates the conceptual data domain; *Subdomain*, indicates subgrouping of the conceptual data domain; *Subdomain category*, indicates subgrouping of the conceptual data subdomain; *Item*, indicates the name of the data item; *Item definition*, describes the data item; *Item conformance*, specifies the conformance of the data item (also known as collection status); *Value sets*, indicates the value sets to be used to specify the item; *Value set reference (Dictionaries or Standards)*, indicates the website hosting the reference database or vocabulary for value sets; *OSIRIS*, indicates, if present, the corresponding data item in the OSIRIS data model; *mCODE*, indicates, if present, the corresponding data item in the mCODE data model; *ICGC-ARGO*, indicates, if present, the corresponding data item in the ICGC-ARGO data model; *GDC*, indicates, if present, the corresponding data item in the GDC data model.

The 1+MG-MDC is also maintained in Zenodo (https://doi.org/10.5281/zenodo.8239363).

## Acknowledgments

The Beyond 1 Million Genomes (B1MG) project has received funding from the European Union Horizon 2020 research and innovation programme under grant agreement No 951724 -B1MG. Genomic Data Infrastructure (GDI) receives funding from the European Union Digital Europe programme under grant agreement number No 101081813 -GDI. This work was also supported by the CCM 2021 “Italian Genomics Strategy: establishment of a steering committee to support the European initiative 1+Million Genomes (1+MG) and Beyond 1+MG (B1MG) and the Inter-institutional Coordination for Genomics in Public Health” project.

---

**Supplementary Table 1.** List of metathesauri and domain-specific dictionaries.

|  |  |  |
| --- | --- | --- |
| International Classification of Diseases for Oncology, Third Edition) (ICDO3, often spelled ICD-O-3) | International double classification system to code neoplastic diseases by specifying both their morphology and anatomical location (topology) | https://www.who.int/standards/classifications/other-classifications/international-classification-of-diseases-for-oncology |
| International Classification of Diseases | Medical classification list curated by the World Health Organization (WHO), containing codes for diseases, signs and symptoms, abnormal findings, complaints, social circumstances, and external causes of injury or diseases | https://icd.who.int/en |
| Systematized Nomenclature of Medicine - Clinical Terms (SNOMED-CT) | Comprehensive, multilingual health terminology concerning both human and non-human medicine and codifying clinical findings, procedures and observable entities | https://www.snomed.org/# |
| Anatomical Therapeutic Chemical (ATC) Classification | Classification of active substances, grouped according to the organ or system on which they act and their therapeutic, pharmacological, and chemical properties | https://www.whocc.no/atc/structure_and_principles/ |
| HemOnc | Medical wiki of interventions, regimens, and general information relevant to the hematology and oncology fields | https://hemonc.org/wiki/Main_Page |
| Surveillance, Epidemiology, and End Results (SEER*Rx) | Interactive antineoplastic drugs database, containing both drugs and regimens | https://seer.cancer.gov/tools/seerrx/ |
| RxNorm | The normalized naming system for generic and branded drugs with links to existing vocabularies, curated by The National Library of Medicine (NLM) | https://www.nlm.nih.gov/research/umls/rxnorm/index.html |
| Unified Code for Units of Measure (UCUM) | All units of measures (used in international science, engineering, and business) | https://ucum.org/trac |
| Logical Observation Identifiers Names and Codes (LOINC) | Set of universal names and ID codes for identifying laboratory and clinical test results | https://loinc.org/ |

